# An open-source pipeline for calcium imaging and all-optical physiology in human stem cell-derived neurons

**DOI:** 10.1101/2025.07.21.664988

**Authors:** Wardiya Afshar-Saber, Federico M. Gasparoli, Ziqin Yang, Nicole A. Teaney, Lahin Lalani, Gayathri Srinivasan, Dosh Whye, Elizabeth D. Buttermore, Kellen D. Winden, Cidi Chen, Mustafa Sahin

## Abstract

High-throughput profiling of neuronal activity at single-cell resolution is essential for advancing our understanding of brain function, enabling large-scale functional screens, and modeling neurological disorders. However, existing approaches are limited by scalability, manual data processing, and variability, thus restricting their ability to detect disease-associated phenotypes. Here, we present a scalable, open-source platform that integrates optogenetic stimulation, calcium imaging, automated data acquisition, and a fully integrated analysis pipeline. By combining spontaneous and evoked activity profiling, the system enables robust quantification of dynamic neuronal responses across hundreds of stem cell-derived human neurons and multiple timepoints, facilitating phenotyping at both cellular and network levels. We validated the platform by recapitulating established activity phenotypes in neurodevelopmental disorders including CDKL5 Deficiency Disorder and SSADH deficiency. In addition, we generated CRISPR-Cas9 knock-in human induced pluripotent stem cell (hiPSC) lines stably expressing the genetically encoded calcium indicator GCaMP6s to model network dysfunction in Tuberous Sclerosis Complex (TSC). Using this system, we further demonstrated functional rescue of the altered neuronal activity observed in the TSC following pharmacological intervention. By linking single-cell dynamics to population-level phenotypes, this framework provides a powerful and broadly applicable tool for disease modeling, mechanistic studies, and therapeutic screening across a range of neurological disorders.

## Introduction

High-throughput, single-cell resolution profiling of neuronal activity is essential for elucidating the mechanisms underlying circuit function, modeling human neurological diseases, and developing targeted therapeutics^1,2^. Traditional electrophysiological techniques such as patch-clamp recording and multi-electrode arrays (MEAs) remain gold standards for functional interrogation of neuronal networks, but their limited scalability, throughput, and spatial resolution constrain their application for large-scale studies of heterogeneous neural populations, particularly in patient-derived stem cell models^2–4^. Genetically encoded calcium indicators (GECIs) have transformed functional imaging of neuronal activity, enabling non-invasive recordings of large neuronal populations with single-cell resolution^5,6^. When combined with optogenetic actuators^7^, these technologies enable all-optical approaches for dissecting network connectivity and synaptic function offering unparalleled opportunities for studying neural circuit dynamics in both health and disease^8,9^.

Several neurodevelopmental disorders such as Tuberous Sclerosis Complex (TSC), CDKL5 Deficiency Disorder (CDD), and Succinic Semialdehyde Dehydrogenase Deficiency (SSADHD) are characterized by alterations in excitability and synchronization that contribute to the pathogenesis of epilepsy^10–12^. Human induced pluripotent stem cell (hiPSC)-derived neuronal models provide a unique opportunity to dissect the altered neuronal activity in human neurons^13–15^; however, scalable tools for quantifying network and single-cell level activities across large populations remain limited. Additionally, most applications of GECIs in human stem cell-derived neuronal models still rely on viral transduction of postmitotic neurons.

While viral delivery is efficient and flexible, it can result in variability in efficiency, heterogeneous expression levels across cells and experiments^16,17^. In contrast, stable genomic integration of GECIs into safe-harbor loci via genome editing offers several advantages, including earlier expression during neural differentiation, enabling longitudinal imaging of circuit maturation and chronic pharmacological interventions^18,19^, while ensuring consistency across hiPSC batches and differentiation protocols^20^. This strategy reduces experimental variability that frequently constrains the detection of subtle, disease-relevant phenotypes in patient-derived models.

Beyond the technical aspects of indicator expression, the lack of robust, scalable acquisition and analysis pipelines remains a major barrier to the routine implementation of all-optical functional assays in stem cell-derived neuronal models. Calcium imaging datasets pose distinct challenges for reliable cell segmentation, extraction of single-cell activity traces, and quantification of network-level dynamics^21^. Although several software offer powerful analysis features, they are often limited by their reliance on manual or semi-automated segmentation^22,23^, are optimized for *in vivo* applications with the lack essential features such as plate mapping^21^ or limited adaptability to *in vitro* all-optical physiology^24^. Fully automated acquisition and analysis pipelines that reduce manual input while enabling high-content, single-cell resolution are essential to realizing the full potential of these approaches for disease modeling and drug discovery.

Here, we present a fully integrated experimental and analytical framework for scalable, high-content calcium imaging in human hiPSC-derived neuronal models. In addition to the commonly used viral transduction method, we knocked in GCaMP6s at the AAVS1 safe-harbor locus using CRISPR-Cas9 and generated hiPSC lines with stable GECI expression. This is combined with a modular, open-source acquisition platform enabling optogenetic stimulation, real-time deep learning-based cell segmentation, and single-cell calcium dynamics quantification via our *PlateViewer* analysis interface. We demonstrated the potential of this approach across multiple differentiation methods and neurodevelopmental models, including TSC, CDD and SSADHD, which share epilepsy as a core clinical feature. We further validated its suitability for pharmacological screening by applying potassium channel modulators to normalize hyperexcitability in *TSC2*-deficient neurons. This integrated platform provides a scalable, reproducible solution for high-throughput functional screening and mechanistic investigation of circuit-level phenotypes in human disease models, with broad applicability for both basic neuroscience and therapeutic development.

## Results

### Generation of stem cell-derived human neurons for calcium imaging and all-optical assay

We generated excitatory neurons using two approaches. On one hand we transduced hiPSCs with a lentiviral vector encoding human NGN2 (hNGN2) under a tetracycline-inducible promoter, alongside a puromycin resistance gene for selection and enrichment (Fig. 1A, 1B)^25^. On the other hand, we formed and dissociated 3D cortical organoids differentiated from hiPSCs at 36 days *in vitro* (DIV) (Fig. 1A)^26^. To support neuronal maturation of both types of differentiating neurons, we co-cultured them with hiPSC-derived astrocytes for ∼50 days. On DIV21 of the hNGN2 differentiation and DIV50 of the organoid-based differentiation, we transduced the co-culture of neurons and astrocytes (iNs) with lentiviral constructs encoding either the GFP-based calcium indicator GCaMP6s or the red-shifted calcium indicator jRCaMP1b and the optogenetic actuator CheRiff, both driven by the human synapsin promoter (pLV-hSynGCaMP6s, pLV-hSyn-CheRiff-eGFP and pLV-hSyn-jRCaMP1b-mRuby) (Fig. 1A). One week post transduction, we observed robust, consistent expression of GFP and mRuby using both differentiation methods, confirming successful transduction of these reporters (Fig. 1C).

**Figure 1.**
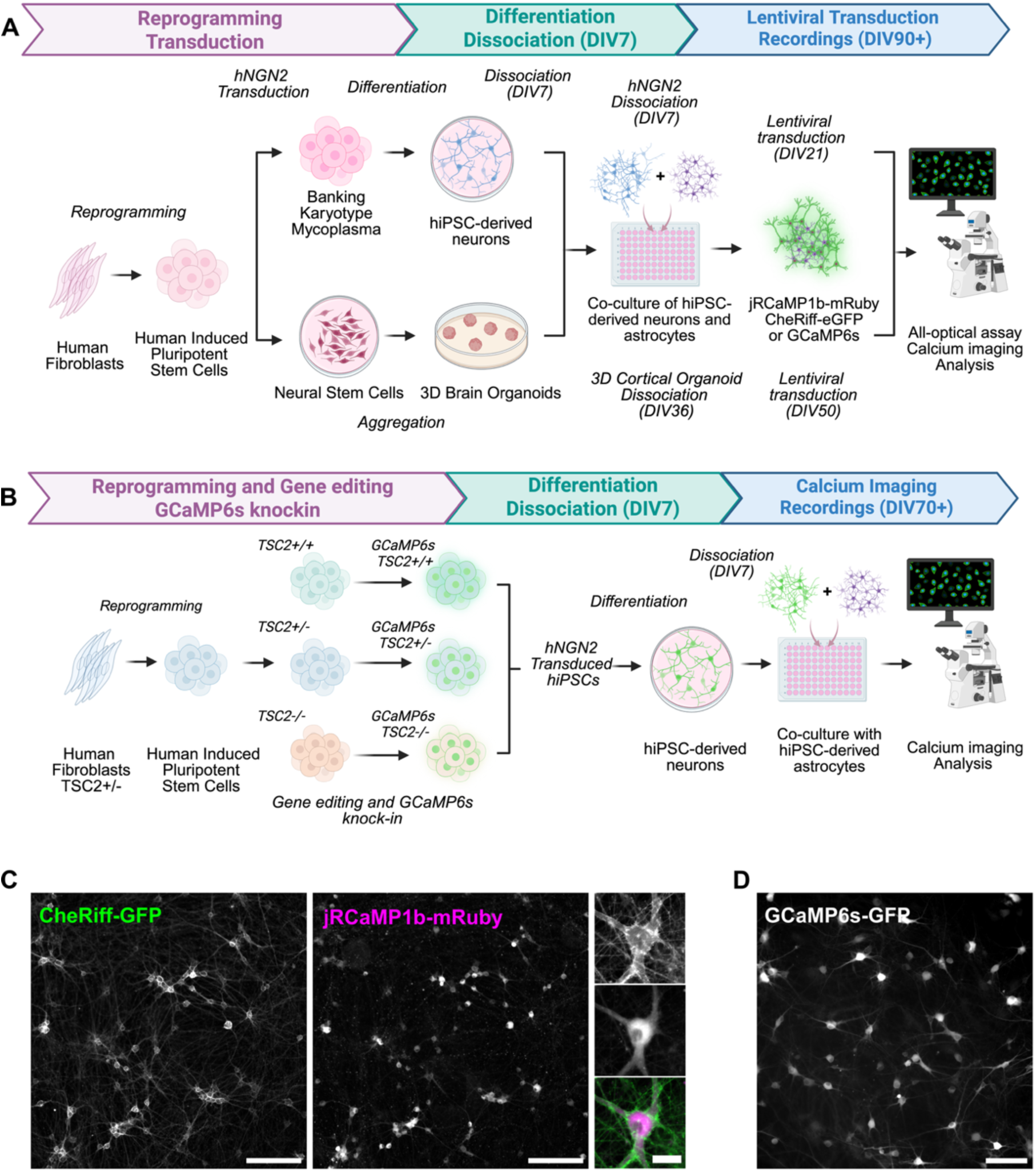
Generation of stem cell-derived neurons for calcium imaging and all-optical physiology. (A) Experimental timeline starting with the reprogramming of human fibroblasts into hiPSCs, followed either by transduction with hNGN2, differentiation and co-culture with hiPSC-derived astrocytes or generation of neural stem cells, 3D cortical organoids followed by dissociation and co-culture with hiPSC-derived astrocytes. Transduction of the calcium indicators for calcium imaging or all-optical assay. (B) Experimental timeline starting with the reprograming of human fibroblasts from TSC patients into hiPSCs, followed by gene editing to generate the full allelic series and CRISPR-mediated GCaMP6s knock-in in the safe harbor locus. After banking, transduction with hNGN2, differentiation and co-culture with hiPSC-derived astrocytes for calcium imaging. (C) Expression of CheRiff-GFP and jRCaMP1b-mRuby in a co-culture of hiPSC-derived neurons and astrocytes at DIV 30, scale bars 200µm and 30µm. (D) Expression of GCaMP6s-GFP in a co-culture of hiPSC-derived neurons and astrocytes at DIV 30, scale bar 100µm. (Created in BioRender. Saber, W. (2025) https://BioRender.com/1is3s0w)

While viral transduction is commonly employed to deliver GECIs into postmitotic neurons, this approach can introduce variability between differentiation batches, posing a challenge for reproducibility in large-scale studies^17^. To overcome these limitations, we also constitutively expressed the GECI *GCaMP6s*-GFP in the AAVS1 safe harbor locus of three hiPSCs lines via CRISPR-Cas9 homologous recombination^18^ (Fig. 1B, Fig. S1A). The first line was derived from a patient with TSC due to an 18bp in-frame deletion in exon 40 of *TSC2* (c.5238_5255del p.H1746_R1751del, *TSC2*^+/−^). The second line was derived from the patient line and had a frameshift mutation induced in the second allele of *TSC2* using TALEN technology (*TSC2*^−/−^), modeling loss-of-function that is observed in TSC. Finally, we generated a third isogenic control hiPSC line (*TSC2*^+/+^) using CRISPR-Cas9^27^ (Table 1). We confirmed successful knock-in clones with the GCaMP6s expression cassette at the AAVS1 (Fig. S1B-C), verified the maintenance of pluripotency by analyzing the expression of pluripotency markers SSEA4, OCT4, Nanog, and TRA-1– 60 and did not detect any karyotypic abnormalities (Fig. S1D-E, Table S2). Finally, the detection of GFP signal in differentiated neurons demonstrated both robust GCaMP6s expression and accurate integration into the safe harbor locus (Fig. S1F).

**Table 1:** Summary table of the hiPSC lines used in this study.

| Neurodevelopmental Disorder | Line name | Sex | Genotype |
| --- | --- | --- | --- |
| Tuberous Sclerosis Complex (TSC) | LAM77 line-TSC2<sup>+/-</sup> | F | c.5238_5255del p.H1746_R1751del (heterozygous) |
|  | LAM77 line-TSC2<sup>-/-</sup> | F | TALEN-LOF TSC2<sup>-/-</sup> homozygous |
|  | LAM77 line-TSC2<sup>+/+</sup> | F | CRISPR-Cas9 TSC2<sup>+/+</sup> homozygous |
| CDKL5 Deficiency (CDD) | HNDS0083-01#B (BCHi005-A) | F | c.1648C>T, p.Arg550* (heterozygous) |
|  | Sex matched parental control | F | c.1648C (homozygous) |
| SSADH Deficiency (SSADHD) | HNDS0005-01#B (BCHi007-A) | F | ALDH5A1<sup>-/-</sup> c.1226G > A / c.1226G > A; p.Gly409Asp |
|  | Sex matched parental control | F | ALDH5A1<sup>+/-</sup> c.1226G / c.1226G > A; p.Gly409Asp |

### Modular acquisition platform for calcium imaging and optogenetics

We developed an open-source acquisition platform incorporating four key features: (1) optogenetic stimulation control, (2) a customizable multi-dimensional acquisition (MDA) interface, (3) real-time segmentation with *Cellpose*^28,29^, and (4) Slackbot-based remote acquisition control (Fig. 2). This platform allowed us to perform large-scale, automated imaging experiments across multi-well plates with high reproducibility and minimal manual intervention (https://github.com/fdrgsp/micromanager-gui).

**Figure 2.**
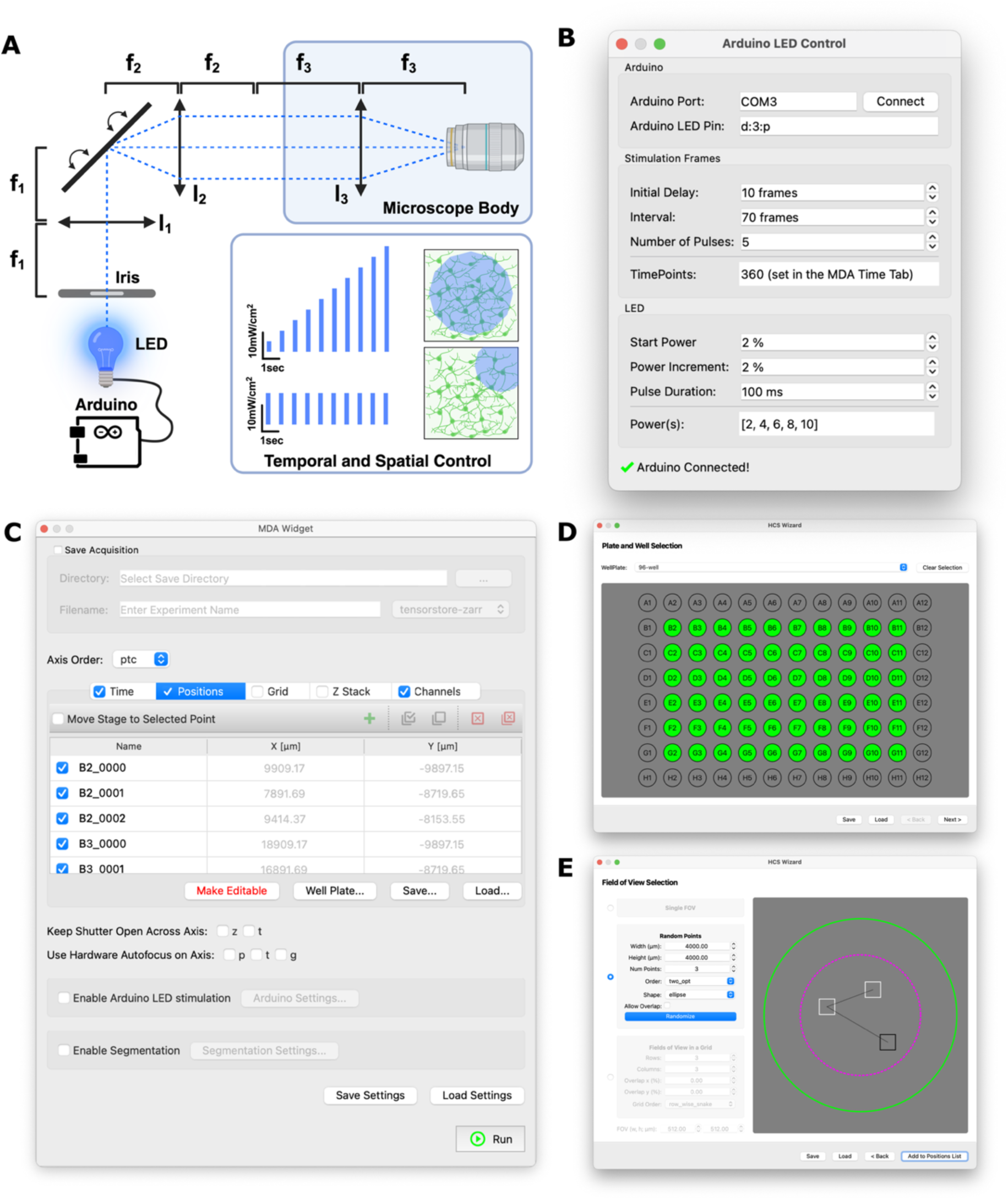
Modular acquisition platform for calcium imaging and all-optical physiology. (A) Microscopy path including an iris for the blue LED stimulation, 20X 0.75NA objective, Arduino for temporal control and iris for spatial control (B) Arduino-based control module for LED pulsing, enabling programmable control of pulse timing, duration, and intensity (C-E) Acquisition software with (C) multi-dimensional acquisition to support key imaging modes including multi-channel, time-lapse, Z-stack, multi-position, and grid acquisitions tailored to calcium imaging (D-E) multi-well plate selection and FOVs selection.

We built our platform on the Micro-Manager ecosystem^30^, leveraging its broad hardware compatibility. While Python offers a widely used and flexible platform for developing custom GUIs, analysis workflows and hardware control, customizing such features within Micro-Manager’s Java-based architecture remains challenging. To address this, we developed a modular graphical user interface (GUI) using the *pymmcore-plus* library (https://pymmcore-plus.github.io/pymmcore-plus), which provides a Python-native interface to the Micro-Manager C++ core. This allowed for rapid customization, seamless integration with modern analysis tools, and direct control of acquisition workflows. Using *pymmcore-plus*, along with *pymmcore-widgets* and *useq-schema* (https://pymmcore-plus.github.io/pymmcore-plus), we developed *micromanager-gui*, an extensible GUI tailored for calcium imaging and optogenetics. The software remains fully compatible with standard Micro-Manager configuration files, allowing users to adopt the platform without modifying existing microscope setups (https://github.com/fdrgsp/micromanager-gui).

To support precise optogenetic stimulation, we developed a dedicated Arduino-based control module for LED pulsing, enabling programmable control of pulse timing, duration, and intensity^8^ (Fig. 2A, 2B). Spatial targeting was achieved using a diaphragm at the sample plane and a kinematically mounted mirror at the back focal plane, allowing us to restrict stimulation to specific subregions of the imaging field (e.g., one corner of a well; Fig. 2A).

For automated data acquisition, we extended the *MDAWidget* to support key imaging modes including multi-channel, time-lapse, Z-stack, multi-position, and grid acquisitions tailored to calcium imaging (Fig. 2C). The widget allowed acquisition settings to be saved and reloaded, ensuring reproducibility across experiments and users. Integration with hardware autofocus (Nikon PFS) maintained stable focus across wells during long-term imaging sessions. To facilitate high-content imaging, we incorporated the *HCSWizard,* a widget that enables plate calibration (which can be saved and reloaded) and flexible field-of-view (FOV) selection across standard plate formats (e.g., 96- or 384-well plates) (Fig. 2D). Users could select center-of-well, reproducible random, or grid-based FOVs, providing flexibility to adapt acquisition strategies to different experimental needs while maintaining consistency and reducing selection bias (Fig. 2E). Additionally, we adopted OME-Zarr^31^ as the standard output format, leveraging Google TensorStore (https://google.github.io/tensorstore/) for asynchronous, chunked data writing during acquisition. This allowed us to capture large time-series datasets efficiently, without memory constraints, and ensured compatibility with scalable, Python-based analysis pipelines.

To further automate the process, users can enable real-time segmentation using a *Cellpose* model (including custom models), which labels fields of view (FOVs) in parallel without disrupting the recording. Additionally, we integrated the Slack API and Slackbot functionality to enable remote control and monitoring of recording progress, enhancing both efficiency and productivity.

### Integrated data exploration and analysis via PlateViewer

We developed *PlateViewer*, an interactive graphical interface integrated into the Micro-Manager GUI, to streamline exploration and analysis of calcium imaging datasets. *PlateViewer* enables visualization, segmentation, and quantitative analysis of multi-well imaging experiments within an intuitive environment (https://github.com/fdrgsp/micromanager-gui). Upon loading a dataset, *PlateViewer* renders an interactive plate layout, allowing users to navigate individual wells and rapidly access associated fields of view (FOVs). For each FOV, the full time series is displayed, enabling direct inspection of spatiotemporal calcium dynamics at single-field resolution. To enable single-cell analysis, we integrated a *Cellpose*-based segmentation interface capable of processing all or selected wells and FOVs^28^. We initially applied the built-in Cyto3 model (Fig. 3A) and subsequently improved its segmentation accuracy by training with labelled images (Fig. 3B). The resulting masks were stored alongside the dataset and served as the basis for subsequent single-cell calcium analyses (Fig. 3C–G). Users can flexibly select subsets of wells or FOVs for processing, facilitating targeted re-analyses and rapid iteration. Following segmentation, *PlateViewer* executes an automated calcium imaging analysis pipeline that extracts raw and normalized calcium traces (ΔF/F or deconvolved) for each segmented neuron and detects calcium transients (Fig. 3C). Detected events are visualized as raster plots in the program, where each row represents an individual neuron, with bar color indicating event amplitude (Fig. 3D). Functional connectivity metrics, including neuronal synchrony (Fig. 3E) and pairwise correlation (Fig. 3F,G), are computed for each FOV. In addition, *PlateViewer* quantifies core activity features such as peak amplitude, event frequency, inter-event interval, and cell size (Fig. 3H-J, https://github.com/fdrgsp/micromanager-gui). As a proof-of-concept, we pharmacologically modulated neuronal activity using CNQX and D-APV, AMPA and NMDA receptor antagonists respectively^25,32^, and caffeine, an adenosine receptor antagonist that enhances network excitability^8,33^. Our acquisition and analysis pipeline reliably identified ROIs across fields of view and quantified fluorescence dynamics in response to these manipulations (Fig. 3H-3J). Caffeine treatment significantly increased calcium event frequency compared to baseline, consistent with its excitatory effects (Fig. 3H). In contrast, synaptic blockade with CNQX/D-APV led to a marked reduction in event frequency (Fig. 3H), a significant decrease in event amplitude (Fig. 3I), and a lower proportion of active neurons (Fig. 3J), as expected with effective suppression of excitatory synaptic transmission. These results demonstrate the sensitivity and robustness of our platform to detect both increases and decreases in network activity in response to pharmacological interventions.

**Figure 3.**
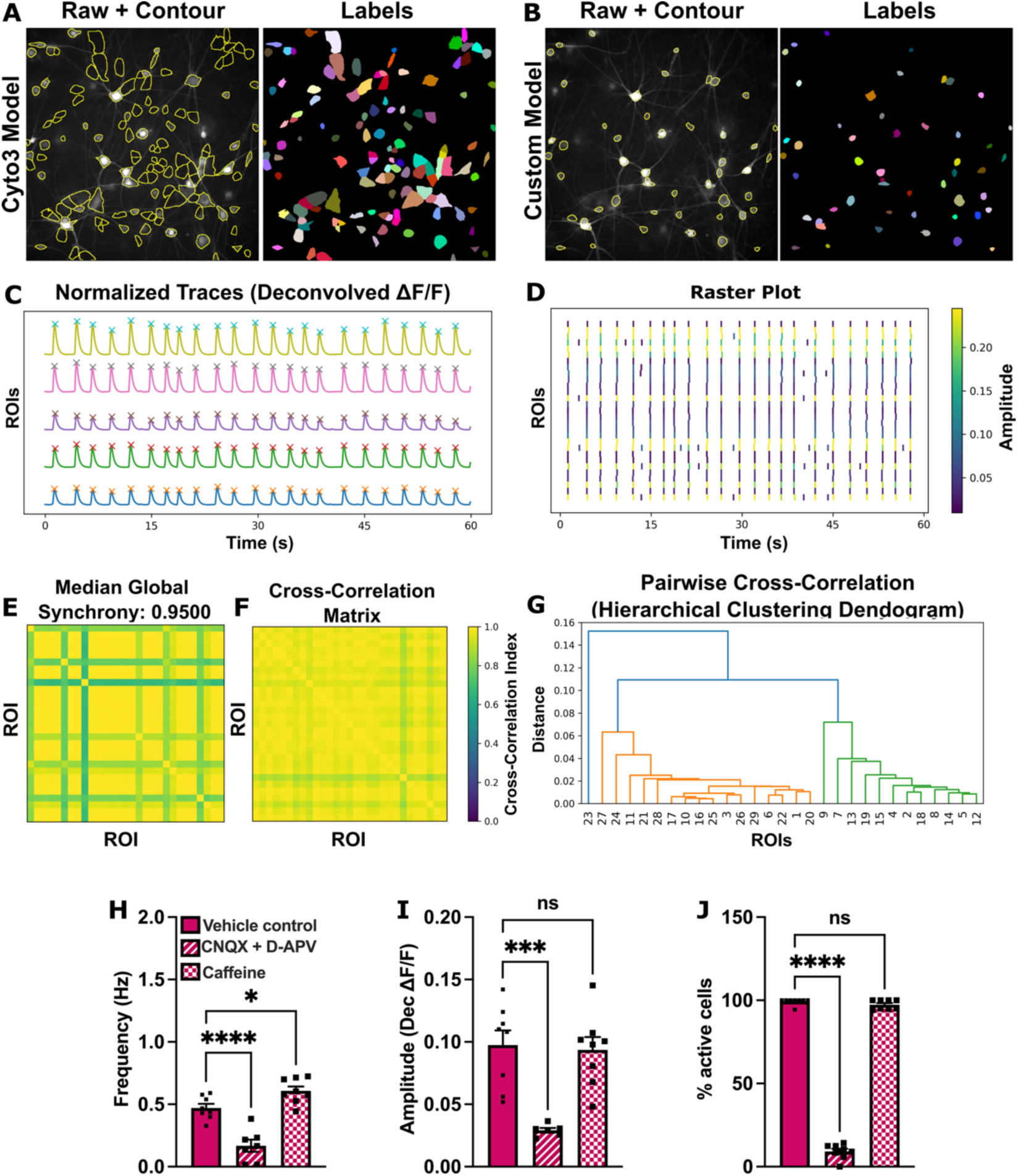
Automated segmentation and analysis pipeline for spontaneous activity. (A–B) Automated segmentation using (A) Cyto3Model and (B) a custom model fine-tuned for improved detection of hiPSC-derived neurons. (C–G) Calcium imaging analysis pipeline: (C) Extraction of fluorescence traces from hSyn-jRCaMP1b-mRuby followed by peak detection (crosses). (D) Raster plots showing calcium transients for individual neurons (ROIs), with color-coded amplitudes. (E) Median global synchrony heatmap representing pairwise correlations across ROIs in each field of view (FOV). (F) Pairwise correlation matrix showing functional connectivity between neurons. (G) Hierarchical clustering dendrogram based on pairwise calcium trace similarity. (H–J) Following acute treatment with CNQX (25 µM) and D-APV (5 µM) or caffeine (5mM), we quantified (H) event frequency (Hz), (I) calcium event amplitude, and (J) percentage of active neurons. VC: 181 neurons; CNQX and D-APV: 161 neurons; Caffeine: 176 neurons; n = 6–8 FOVs per condition (mean ± S.E.M; ****p < 0.0001; ***p < 0.001; *p < 0.005; ns = non-significant). One-way ANOVA followed by Dunnett’s multiple comparisons test (H) F (2, 20) = 32.81, ****p < 0.0001 and *p = 0.0342; (I) F (2, 19) = 13.46, p = 0.0003 and (J) F (2, 21) = 1936, ****p < 0.0001 and ^ns^p = 0.4075.

### Quantification of Optically Evoked Activity

We further developed an evoked activity analysis mode within *PlateViewer* to quantify neuronal responses to optogenetic stimulation (Fig. 4). Neurons co-expressing CheRiff and jRCaMP1b (Fig. 1A, 1C) were stimulated with 100ms blue light pulses applied in 1% increments from 1% to 10% intensity (10 pulses; ∼4mW/cm² to ∼100mW/cm²). *PlateViewer* extracted fluorescence signals from individual ROIs, detected stimulus-evoked peaks (Fig. 4B,C), and quantified response amplitudes as a function of irradiance (Fig. 4D). Neuronal responses were detectable at the lowest intensity (∼4 mW/cm²) and saturated at ∼48mW/cm². To further characterize stimulus-response relationships, we varied pulse duration (1, 10, 50, and 100ms). Short pulses (1ms and 10ms) elicited responses above baseline noise, while longer pulses (50ms and 100ms) produced more robust and quantifiable activity, plateauing at ∼48mW/cm² (Fig. 4D).

**Figure 4.**
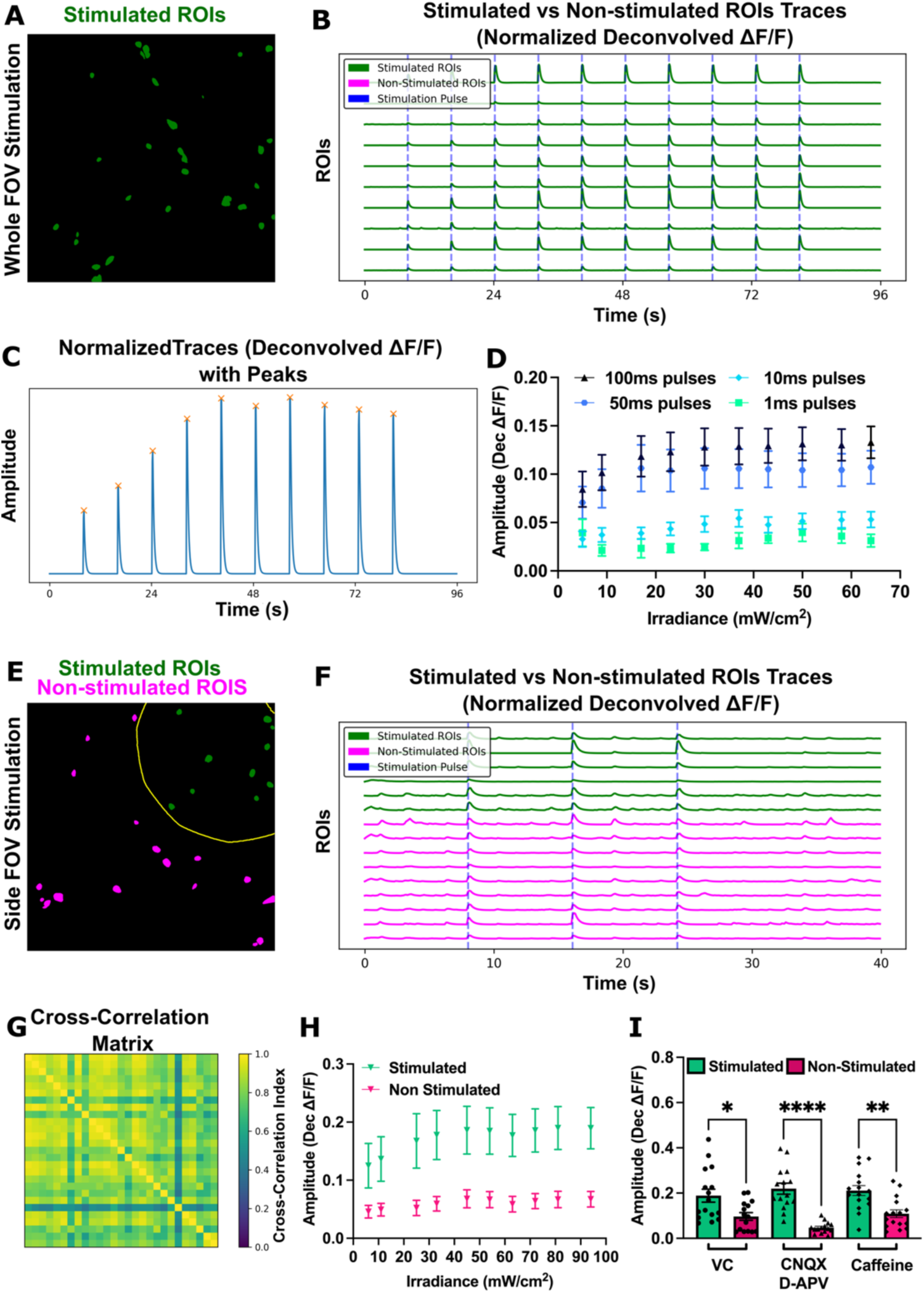
Evoked Activity Analysis Pipeline. (A) Segmentation mask using our custom model showing all stimulated neurons in green; non-stimulated cells in magenta (none in this experiment). (B) Calcium traces from 10 randomly selected ROIs, with blue dashes indicating stimulation pulses. (C–E) Partial field stimulation: segmentation mask highlighting stimulated (green) and non-stimulated (magenta) neurons within the yellow stimulation boundary. (F) Calcium traces from 10 random ROIs, color-coded by stimulation status; blue dashes indicate timing of blue-light pulses. (G) Pairwise cross-correlation matrix illustrating functional connectivity across all ROIs, with color scale indicating correlation strength. (H) Calcium event amplitudes in stimulated (green) vs. non-stimulated (magenta) neurons across increasing light intensities. (I) Calcium event amplitudes in stimulated and non-stimulated neurons following acute treatment with synaptic blockers (CNQX: 25 µM, D-APV: 5 µM) or caffeine (5 nM) One-way ANOVA followed by Tukey’s multiple comparisons test n = 6-8 FOVs, 2 independent differentiations; F (5, 87) = 11.64 (mean ± S.E.M; ****p < 0.0001; **p < 0.01; *p < 0.05).

To enable spatially resolved analyses, *PlateViewer* incorporates stimulation masks that automatically classify ROIs based on their position relative to the stimulated region (Fig. 2A, 4E). Subregion-specific stimulation allowed segregation of stimulated and non-stimulated neuron populations within the same FOV, enabling direct comparison of local and network-wide activity following stimulation (3 pulses, 100ms, 30% intensity) (Fig. 4F). Finally, *PlateViewer* supports cross-correlation analyses to assess network-level responses and detect changes in synaptic transmission (Fig. 4G).

As proof-of-concept, we optically stimulated a subregion of wild-type hiPSC-derived neurons co-expressing CheRiff and jRCaMP1b (10 pulses, 100ms, ∼4–100mW/cm²), and recorded activity in the entire FOV (Fig. 4H). Stimulated neurons consistently exhibited higher response amplitudes across all intensity levels, plateauing at ∼48mW/cm². To demonstrate the sensitivity of the system to pharmacologically induced activity changes, we modulated synaptic activity as previously shown in primary neurons^8^, using synaptic blockers and caffeine. As anticipated, synaptic blockers significantly reduced activity in non-stimulated neurons compared to vehicle control, while caffeine increased non-stimulated neuronal responses, confirming the sensitivity of the acquisition and analysis pipeline to detect network-level changes in activity (Fig. 4I).

### Application in disease modelling

To validate the robustness and versatility of our acquisition and analysis pipeline across distinct neurodevelopmental disorders, we applied it to models of TSC, CDD, and SSADHD using complementary strategies for GECI expression, indicator types, and neuronal differentiation protocols. For TSC, we generated an isogenic allelic series of hiPSC lines with stable GCaMP6s integration into the AAVS1 safe harbor locus in *TSC2*^+/+^, *TSC2*^+/−^, and *TSC2*^−/−^ backgrounds (Table 1, Fig. 1B, 1D; Fig. 5A–5C), ensuring consistent long-term expression from early differentiation through neuronal maturation and chronic treatments. In parallel, for CDD (Table 1, Fig. 5D–5G) and SSADHD models (Table 1, Fig. 5H–5J), we transduced post-mitotic neurons with GCaMP6s or jRCaMP1b via lentiviral delivery (Fig. 1A, 1C, 1D). Neuronal differentiation employed an accelerated cortical differentiation protocol for CDKL5 and hNGN2-mediated differentiation protocol for TSC and SSADH deficiency, enabling assessment of disease-associated phenotypes across multiple models.

**Figure 5.**
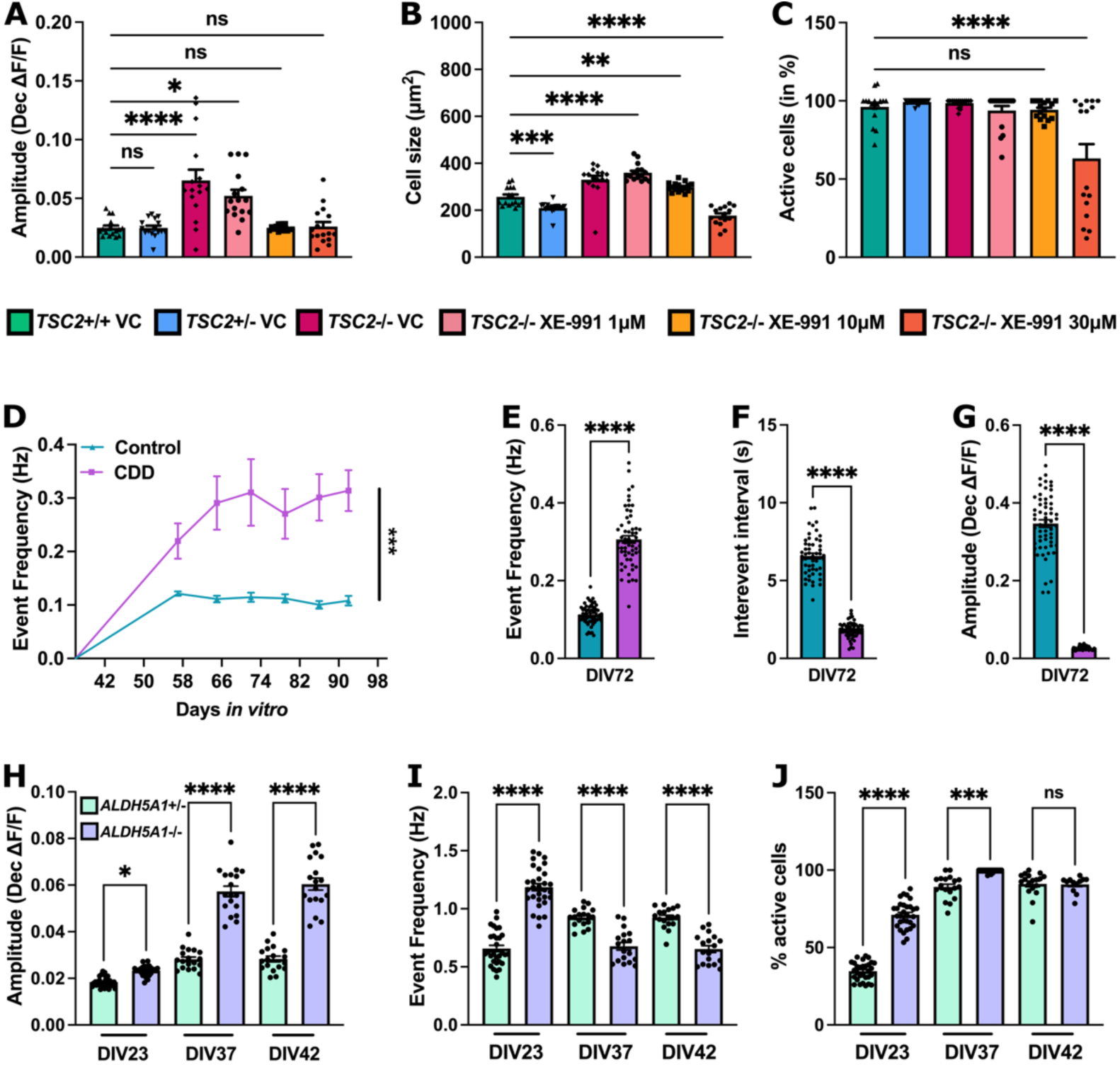
Application in disease models. (A-C) Quantification of the activity in TSC2^+/+^, TSC2^+/-^, and TSC2^-/-^ hiPSC-derived neurons chronically treated with XE-991 at (1, 10 and 30µM) showing (A) increased in amplitude in TSC2-deficient neurons, rescued in the XE-991 treated conditions, (B) cell size showing significantly larger cells TSC2-deficient neurons compared to controls unchanged upon treatment with XE-991 at 1 and 10µM but decreased at the highest concentration (30µM) and (C) percentage of active cells, decreased in the treated condition with 30µM. (A-C) One-way ANOVA followed by Dunnett’s multiple comparison test (A) F (29, 438)=18.69; (B) F (29, 450)=50.69; (C) F (29, 450)=9.391, n=3 independent differentiations, 8FOVs/condition; TSC2^+/+^: 178 ROIs; TSC2^+/-^: 852 ROIs; TSC2^-/-^: 347 ROIs; TSC2^-/-^ 1µM XE-991: 471 ROIs; TSC2^-/-^ 10µM XE-991: 517 ROIs; TSC2^-/-^ 30µM XE-991: 298 ROIs. (D-G) Quantification of the activity in CDD and sex matched parental control with (D) longitudinal quantification of event frequency in hertz showing sustained network hyperexcitability in CDD neurons (E) At DIV72, detailed analysis revealed (F) significantly shorter inter-event intervals and (G) reduced calcium event amplitudes relative to controls. (E-G) Unpaired Welch’s t test (E) F(58,57)=8.369 (I) F(51,57)=6.601 (G) F(58,54)=292.4, n=1 independent differentiation, 60 FOVs/condition with Control: 6620 ROIs; CDD: 1889 ROIs. (H-J) Quantification of the activity in SSADH-deficient neurons ALDH5A1^-/-^ and sex matched parental control ALDH5A1^+/-^ at three timepoints DIV23, DIV37 and DIV42 with (D) quantification of amplitude showing increase in ALDH5A1^-/-^, (I) event frequency showing more frequent events at DIV23 and lower frequency as the network mature at DIV37 and 42 in the SSADH deficient neurons and (J) percentage of active cells increasing over time in both conditions while a significantly greater proportion of SSADH-deficient neurons were active at early developmental stages (DIV23). (H-J) One-way ANOVA followed by Dunnett’s multiple comparison test (H) F (5, 126) = 185.4; (I) F (5, 126) = 65.27; n=3 independent differentiations, 18FOVs/condition; ALDH5A1^+-+^: 978 ROIs; ALDH5A1^-/-^: 944 ROIs (mean ± S.E.M; ****p < 0.0001; **p < 0.01; *p < 0.05; ns=non-significant).

Using our automated pipeline, we recorded spontaneous calcium activity in *TSC2*^-/−^ neurons at DIV42 and observed a significant increase in calcium event amplitude (Fig. 5A) and cell size (Fig. 5B) relative to *TSC2*^+/−^ and *TSC2*^+/+^ controls, consistent with previously reported phenotypes in hiPSC-derived neurons^14^. To probe therapeutic potential, we chronically treated TSC2-deficient neurons with the potassium channel modulator XE-991 at 1, 10, and 30µM from DIV10 to DIV42^34^. We found that 1µM XE-991 had no effect on calcium event amplitude, while 10 and 30µM treatments significantly reduced event amplitude to control levels (Fig. 5A), indicating dose-dependent normalization of neuronal hyperactivity. XE-991 at 1 and 10 µM did not affect cell size; however, 30µM induced significant reductions in both cell size (Fig. 5B) and the proportion of active neurons (Fig. 5C). These findings were further confirmed by visible neuronal deterioration in the calcium imaging recordings. These results demonstrate that targeted potassium channel modulation can rescue hyperexcitability in *TSC2*-deficient neurons at non-toxic concentrations.

Next, we modeled CDD, a neurogenetic disorder characterized by early-onset seizures and disrupted network dynamics. CDKL5-deficient hiPSCs and isogenic control were differentiated using an accelerated cortical differentiation protocol^26^. We dissociated organoids at DIV35, co-cultured neurons with hiPSC-derived astrocytes, transduced with pLV-hSyn-GCaMP6s, and performed longitudinal calcium imaging every 6-8 days from DIV21 to DIV56. Compared to controls, CDKL5-deficient neurons consistently displayed elevated calcium event frequency at all timepoints, indicating sustained network hyperexcitability (Fig. 5D). At DIV36, detailed analysis revealed significantly shorter inter-event intervals and reduced calcium event amplitudes relative to controls, reflecting altered neuronal firing dynamics. These findings are consistent with previously reported functional deficits in CDKL5 deficiency^35^ and underscore the sensitivity of our pipeline for detecting disease-relevant activity changes.

Finally, to model SSADHD, a neurodevelopmental disorder linked to epilepsy, we generated *ALDH5A1*^−/−^ patient-derived hiPSCs and sex matched parental control *ALDH5A1*^+/−^. Neurons were differentiated using hNGN2 overexpression, co-cultured with hiPSC-derived astrocytes and profiled longitudinally at DIV23, DIV37, and DIV42. We observed persistently elevated calcium transient amplitudes in SSADH-deficient neurons at all time points. Early in development (DIV23), calcium event frequency was significantly higher than controls, indicating initial network hyperexcitability. As networks matured, event frequency progressively declined while high-amplitude transients persisted, indicative of emerging hypersynchronous, burst-like activity reminiscent of epileptiform discharges. Additionally, a significantly greater proportion of neurons were active at early developmental stages in SSADH-deficient cultures. Collectively, these dynamic network alterations recapitulate key phenotypes previously reported in hiPSC-derived neuronal models of SSADH deficiency^13^.

Together, these results validate the broad applicability and generalizability of our platform to capture distinct, disease-relevant neuronal phenotypes across multiple genetic neurodevelopmental disorders, enabling mechanistic insights and providing a scalable framework for high-content functional screening and therapeutic development.

## Discussion

The integration of functional imaging with human iPSC-derived neuronal models offers transformative opportunities to dissect the cellular and circuit-level mechanisms underlying neurodevelopmental disorders. Here, we present a fully integrated, scalable, and open-source platform that combines optogenetic stimulation, GECIs, multi-well imaging, and automated single-cell analysis to capture both spontaneous and evoked network activity. Applying this system across three distinct neurodevelopmental disorders (TSC, CDD and SSADHD), we demonstrated its versatility for uncovering disease-relevant phenotypes and its utility as a discovery platform for therapeutic interventions in precision medicine.

Functional calcium imaging offers a unique advantage: longitudinal, non-invasive measurement of large neuronal populations with single-cell resolution. Additionally, by incorporating spatially patterned optogenetic stimulation, our platform further allows the interrogation of synaptic integration and subnetwork dynamics, revealing how altered activity disrupt both intrinsic excitability and circuit-level function^8,9^. Despite the availability of several calcium imaging analysis platforms that provide robust tools for quantification, most face key barriers to scalability and broad applicability. Common bottlenecks include manual or semi-automated cell segmentation workflows, which limit throughput and reproducibility, as well as restricted capabilities for integrating multiple experimental conditions or conducting longitudinal studies *in vitro*^21–23^. These limitations pose particular challenges for large-scale disease modeling and drug screening using human iPSC-derived neurons, where experiments often involve multiple timepoints, genotypes, and pharmacological conditions.

By contrast, a major advance of our platform lies in its automation and scalability. The fully integrated acquisition and analysis pipeline, incorporating real-time deep learning-based segmentation, automated trace extraction, and interactive data visualization via the *PlateViewer* interface, significantly reduces user bias and technical variability, enabling reproducible, large-scale studies across multiple differentiation batches and patient lines making it uniquely suited for scalable, single-cell resolution studies of circuit-level phenotypes in neurodevelopmental disorders. We demonstrated the sensitivity of this system to detect both hyperactive and hypoactive network states in response to pharmacological modulation and to reveal gene-specific and patient-specific functional signatures in CDD^35,36^ and SSADHD^13^ models, highlighting its potential for drug screening.

Additionally, a key technical innovation of our approach is the generation of GCaMP6s knock-in hiPSC lines via CRISPR/Cas9-mediated targeting of the AAVS1 safe harbor locus^18^, ensuring stable, homogeneous, and early expression of GECIs throughout neural differentiation. This strategy enables consistent longitudinal imaging across development, reduces variability across experiments, and facilitates direct comparisons between isogenic control lines and patient-derived models. Using this approach, we validated previously reported network hyperexcitability phenotypes in TSC using multielectrode array (MEA) recordings and demonstrated that pharmacological inhibition of KCNQ channels with XE-991, a compound shown to normalize hyperactivity in TSC models^34^, effectively modulates network excitability in our system.

Finally and importantly, the modular and open-source architecture of this platform fosters rapid dissemination, adaptation, and community-driven development. It provides a flexible framework that can be readily extended to additional disease models, imaging modalities, and perturbations. Future directions include integration of multimodal datasets such as transcriptomics, proteomics, and metabolomics to build comprehensive, multiscale models of disease, as well as incorporation of machine learning approaches for phenotype classification and functional connectivity mapping.

Together, these developments will enhance the platform’s capacity to identify novel subtypes, molecular targets, and biomarkers. In summary, this work addresses a critical gap in the field by enabling scalable, high-content functional phenotyping of human neural circuits. By combining all-optical physiology, automated analysis, and disease modeling in patient-specific iPSC neurons, our platform lays the foundation for mechanistic insights and therapeutic discovery in complex neurological disorders.

## Methods

### Generation of Human Induced Pluripotent Stem Cells (hiPSCs)

Subjects for this study were recruited through Boston Children’s Hospital, and the study protocol was approved by the institutional review board (IRB-P00008224, IRB-P00016119). Informed consent was obtained from all participants and/or their parents, as appropriate. Subject characteristics and reprogramming methods have been previously described^27,37,38^ and are listed in Table 1.

### Generation of knock-in GCaMP6s hiPSCs

Based on the human PPP1R12C genomic assembly GRCh38.p13 on NCBI, gRNAs were designed to target the AAVS1 area in the intron region. sgRNA: GGGGCCACTAGGGACAGGAT (TGG). The AAVS1-Puro-CAG-GCaMP6s plasmid was a gift from Xiaojun Lian (Addgene plasmid # 20896; http://n2t.net/addgene:120896; RRID: Addgene_120896). Briefly, the plasmid targets the AAVS1 safe harbor locus and contains the GCaMP6s gene under the CAG promoter and the puro resistance gene with homology-directed repair.

The GCaMP6s reporter was inserted into the AAVS1 safe locus of the isogenic hiPSC sets using CRISPR/Cas9 techniques. The plasmid was delivered via electroporation using the Invitrogen Neon transfection system (Thermo Fisher, #MPK5000). Briefly, the hiPSCs were digested by 0.75x TrypLE Select (Gibco, #12563029) in dPBS (Gibco, #14190-144) at 37°C for 5 min. 1 million single cells were electroporated with 1.1ul 100µM sgRNA, 2µg AAVS1-GCaMP6s plasmid, and 1.5µl Alt-R® S.p. HiFi Cas9 Nuclease V3 (IDT, #1081061) in the Resuspension R buffer (ThermoFisher, #MPK10096) at 1200V, 30ms, and 1 pulse in a cuvette. The resulting cells were plated onto one well in a Cultrex-coated 6-well plate (ThermoFisher, #140675) in mTeSR plus with 1:10 10x cloneR2 (StemCell Technology, #100-0691) and 1:100 3.55mM Nu7026 (SelleckChem, #S2893) and left at room temperature for an hour before incubating at 37°C and 5%𝐶𝑂_!_. 24 hours later, the medium was changed to fresh mTeSR plus with 10µM Y27632 (Cayman Chemical, 10005583). Afterwards, the medium was changed to mTeSR plus for every other day feeding. When the cells reached 40-60% confluency, 0.6µg/ml puromycin (InvivoGen, ant-pr-1) was added to the mTeSR plus media for selection for 3 days. The culture was then maintained in mTeSR+ with every other day feeding until it reached about 70% confluency. 4000 single cells were seeded per 10cm dish (ThermoFisher, #150464) for colony picking into Cultrex-coated 96-well plates (Corning, #353072). When the visible colonies were formed, 24 clones per genotype were expanded to 24-well plates (ThermoFisher, #142485). Specifically, ⅓ of the cell resuspension was passaged, ⅓ was purified for gDNA, and ⅓ was cryopreserved. AmpliTaq PCR on the extracted gDNA confirmed the homozygous insertion of the calcium reporter with two pairs of primers, Leftarm-Fw and Rightarm-Rv; Intron-Fw2 and Puro-Rv (Table S1). 4-5 clones per genotype with homozygous insertion were cryopreserved. One clone per genotype was further characterized for pluripotency markers, karyotypes, and Sanger sequencing for the insertion site (Table S1). After confirming the quality of the cell lines, the clones were expanded, cryopreserved, and proceeded with downstream experiments.

### Genomic DNA AmpliTaq PCR

After the culture reached 70% confluency, the cells were rinsed with 1 mL dPBS once and pelleted by scraping in 1 mL dPBS. The cell lysate was collected into one 1.5 ml Eppendorf tube and centrifuged at 15000 rpm for 30 seconds. The genomic DNA was purified using the Monarch Spin gDNA Extraction kit (New England Biolabs, #T3010L). Briefly, the cell pellets were resuspended in 100µl cold dPBS. 3µl Protein kinase K and 3µl RNase A were added to the resuspension and mixed thoroughly by brief vortexing. 100µl cell lysis buffer was then added and mixed by vortex. The mixture was incubated at 56°C in a thermal mixer with agitation at 1400 rpm. 400µl gDNA binding buffer was added to the resuspension and mixed thoroughly by pulse-vortexing for 10 seconds. The lysate and buffer mixture were transferred into a gDNA purification column pre-inserted into a collection tube and centrifuged first at 1000 rpm for 3 min and then at 15000 rpm for 1 min. The flow-through and the collection tube were discarded. The column was then washed twice with 500µl gDNA wash buffer, followed by centrifugation at 15000 rpm for 1 min each. The flow-through and the collection tube were discarded again. The column was inserted into a labeled, cleaned Eppendorf tube and incubated at room temperature in 100µl RNA-free water preheated at 60°C for 1 min before being centrifuged again at 15000 rpm for 1 min. The concentration of nucleic acid in the final flow-through was determined using the Tecan Spark10M multimode microplate reader.

The PCR was carried out using the AmpliTaq Gold 360 DNA Polymerase kit (ThermoFisher, #4398881). Briefly, 25µl of AmpliTaq Gold 360 Master Mix, 5µl of GC enhancer, 2µl of primers (1µl forward and 1µl reverse), and 18µl of gDNA and UltraPure™ DNase/RNase-Free Distilled Water (Invitrogen, #10977015) were mixed for one 50µl reaction. Between 50-200ng gDNA was loaded, depending on the concentration, and all samples were loaded with the same weight. The PCR reaction was run on a BioRad C1000 Touch Thermal Cycler (BioRad, #1851148). To confirm the insertion, E-Gel 2% Agarose Gels with SYBR Safe (Invitrogen, #A42135) were used for electrophoresis. PCR products were also sent for Sanger Sequencing to ensure the correct insertion sequences.

### Mycoplasma testing

All cellular cultures were routinely tested for mycoplasma by PCR and only negative samples were used in this study. Media supernatants (with no antibiotics) were collected, centrifuged, and resuspended in a saline buffer. Ten microliters of each sample were used for a PCR with the followings sets of primers: Myco280_CReM (5′-ACACCATGGGAGYTGGTAAT-3′); Myco279_CReM (5′-CTTCWTCGACTTYCAGACCCAAGGCAT-3′) from the Center for Regenerative Medicine: CReM - Boston University and MGSO-5 (5′-TGCACCATCTGTCACTCYGTTAACCTC-3′) and GPO-3 (5′-GGGAGCAAACAGGATTAGATACCCT-3′).

### Immunocytochemistry and imaging

Immunocytochemistry staining was performed to confirm the pluripotency of the generated hiPSC lines. The cells were cultured on poly-D-lysine (Sigma-Aldrich, P6407-5MG) /Laminin (ThermoFisher, #23017-015)-coated 96-well plate (Cellvis, #P96-1.5P) and fixed using 4% paraformaldehyde (PFA; Polysciences, #04018-1) in PBS when they reached ideal confluency. Briefly, an equal amount of 8% PFA was added to each well on top of the media. The plate was incubated in 4% PFA at room temperature for 20 minutes, followed by three rolling washes with 100µl/well PBS. The plate was stored at 4°C if not stained immediately. The blocking buffer was made as follows: 5% Normal Goat Serum (ThermoFisher, #50197Z), 2% Bovine Serum Albumin (Gibco, #15260-037), and 0.1% Triton-X 100 (Sigma-Aldrich, #T9284-100ML) in PBS. The fixed plate was incubated in 100µl/well blocking buffer at room temperature for an hour before 50µl/well primary antibody diluted in blocking buffer was added (Table S2) and stored at 4°C overnight. Approximately 16 hours later, the primary antibodies were removed, and the cells were washed with 100µl/well PBS three times. The cells were then incubated with 50µl/well secondary antibodies (Table S2) diluted in blocking buffer at room temperature for an hour in the dark. After the incubation, the antibodies were removed, and the cells were washed with 100µl/well PBS three times in the dark. After the washes, the cells were incubated in 4µg/ml Hoechst (Invitrogen, #H3569) diluted in UltraPure 𝑑𝑖𝐻_!_𝑂 for 5 minutes, followed by three washes with 100µl/well 𝑑𝑖𝐻_!_𝑂. The stained plates were imaged with the ImageXpress Micro XLS High-Content Imaging System (Molecular Devices) at 20x magnification with a 6-step z-stack.

### Generation of hiPSC-derived neurons (hNGN2)

We used two differentiation methods in this manuscript, the NGN2 overexpression^39^ with minor changes as described in one of our previous studies^13^ and an accelerated cortical organoid differentiation^26^. For the NGN2 differentiation, the vectors were a gift from Kristen Brennand transduced with lentiviral vectors that encode human NGN2 (hNGN2) under the tetracycline promoter, as well as the puromycin (pLV-TetO-hNGN2-Puro, addgene # 79049 ; http://n2t.net/addgene:79049 ; RRID:Addgene_79049) or neomycin (pLV-TetO-hNGN2-Neo, addgene #99378 ; http://n2t.net/addgene:99378 ; RRID:Addgene_99378) resistance gene for selection. hiPSCs were grown under feeder-free conditions, mycoplasma negative, karyotyped after transduction, and no abnormalities were detected. The hNGN2 transduced hiPSCs were dissociated into single cells using Accutase and seeded onto Geltrex-coated plates at a density of 100,000/cm^2^. The next day, hNGN2 expression was induced using doxycycline and selected with puromycin. Growth factors BDNF (10 ng/ml, catalog #450–02; Peprotech), NT3 (10 ng/ml, catalog #450–03; Peprotech), and laminin (0.2 mg/l, catalog #23017–015; Thermo Fisher Scientific) were added in N2 medium for the first two days. Cells were then fed with BDNF (10 ng/ml), NT3 (10 ng/ml), laminin (0.2 mg/l), doxycycline (2 μg/ml), and Ara-C (4uM, catalog #C1768; Sigma-Aldrich) in B27 media and fed every other day until dissociation at DIV7. Cells were then dissociated with papain (catalog #LK003178; Worthington) and DNaseI (catalog #LK003172; Worthington) and replated on 96well plate (Cellvis, #P96-1.5P) or 384 plates Poly-D-Lysine (PDL 0.5 mg/ml; catalog P6407; Sigma Aldrich) and laminin (5 μg/ml; catalog #23017–015; Life Technologies) coated plates at a density of 1300/mm^2^ in co-culture with hiPSC-derived astrocytes at a density of 200/mm^2^ (Astro.4 U; Ncardia).

### Generation of 3D accelerated cortical organoids and 2D dissociated cultures

hiPSC clones that express wild-type CDKL5 (BCHi005-B) and the variant form (BCHi005-A) derived from a female patient with CDKL5 deficiency disorder^38^ were first differentiated to neural stem cells (NSCs) in a monolayer format as outlined in a previous study^26^. A large stock of NSCs was established by freezing the cells in a cryopreservation media containing 90% culture media and 10% DMSO on day 11 of differentiation^36^. NSCs from this bank were thawed, expanded until day 14 of differentiation and were aggregated at 5 million cells/well of a 6 well plate and were differentiated to cortical organoids as previously described^26^. For dissociating the organoids and plating them for calcium imaging, first, 96 well glass bottom plates (Cellvis # P96-1.5P) were coated with 25µg/mL poly-L-ornithine (PLO; Sigma Aldrich #P4957) in borate buffer (Thermofisher #28341), overnight at 37°C. The PLO solution was aspirated and the wells were next coated with 5µg/mL laminin solution (Thermofisher #A29248) in DPBS overnight at 37°C. Laminin solution was aspirated just before use. One day prior to dissociating the organoids, astrocytes (NCardia Astro.4U) were plated onto the inner 60 wells of the poly-L-Ornithine/laminin-coated 96 well glass bottom plates at 6000 cells/well in astrocyte media (Sciencell #1801) with 10µM rock inhibitor (Cayman Chemical #10005583). The outer wells were filled with sterile water to lessen evaporation of media from the wells on the edge. On day 36 of differentiation, organoids were dissociated into single cells by incubating them in Accumax (Innovative Cell Technologies # AM105.500) for approximately 2 hours at room temperature followed by quenching with 10% knockout serum replacement (Invitrogen #10828028) in DMEM. The organoids were gently triturated and the cells were passed through a 40µm cell strainer. Cells were counted and plated on astrocytes in the 96 well plates at 40,000 cells/well in phenol red-free BrainPhys media (Stemcell Technologies # 05791) supplemented with SM1 supplement (Stemcell Technologies # 05711), GlutaMax (Thermofisher #35050061) and BDNF (Peprotech 450-02), NGF (Peprotech #450-01), NT3 (Peprotech #450-03), laminin, 10µM rock inhibitor and CEPT (Tocris 7991). Cells were plated in half the final volume and 24 hours after plating the rest of the media without rock inhibitor and CEPT was added. Half media changes were performed using a P300 Integra multichannel pipette twice a week.

### Calcium imaging: Spontaneous and Evoked Activity

Cultures of iNs and hiPSC-derived astrocytes were transduced with lentiviral particles (4hours incubation) pLV-hSyn-jRCaMP1b at DIV 21 for hNGN2 and DIV50 for the dissociated cortical organoids and recordings weekly starting one week post transduction. Viral particles were prepared by the Viral Core at Boston Children’s Hospital. Imaging was performed on a Nikon Ti2-E epifluorescence microscope (https://tinyurl.com/mic-all-optical) equipped with a Nikon 20×/0.75 NA air objective and a PCO Edge 4.2 USB sCMOS camera. Samples were maintained at 37°C with 95% humidity and 5% CO₂ using a stage-top incubator (Okolab, model H01-T-UNIT-LB-Plus). The change in fluorescence from jRCaMP1b was recorded at 10Hz, binning 2, under continuous green illumination at 550nm through a Chroma filter set: excitation filter ET570/20x, dichroic mirror T585lpxr, longpass emission filter ET590lp. The change in fluorescence from GCaMP6s was recorded at 10 or 20Hz, binning 2, under continuous blue illumination at 470 nm through a Chroma filter set: excitation filter ET470/40x, dichroic mirror T495lpxr, emission filter ET525/50m). For the evoked activity experiments, the blue illumination for the stimulation of the light-activated ion channel CheRiff, was provided by a blue light LED at 455 nm (Thorlabs M455L3 mounted LED with LEDD1B Driver) through a Chroma filter set: excitation filter ET470/40x, dichroic mirror T505lpxr (installed on the second filter turret of the microscope). A lever-actuated iris diaphragm (Thorlabs CP20S) was installed after the blue LED in a plane conjugated with the focal plane of the microscope objective. In this way, within the field of view, different sizes of the blue illumination could be selected. An elliptical mirror (Thorlabs) mounted on a Thorlabs kinematic mount (KCB1EC/M) was placed in a plane conjugated with the back focal plane of the objective so that steering it would allow for positioning the blue illumination in different part of the FOV. Iris + kinematic most allows for full control of illumination size and position. This was achieved by relaying the iris plane and the mirror plane with a set of Thorlabs lenses (l1=AC254-100-A, l2=AC254-100-A and l3=AC254-150-A in Fig. 2) in a 4f configuration.

### Multi-Dimensional Acquisition (MDA) Interface

To support scalable, open-source data acquisition for calcium imaging and optogenetic stimulation across multi-well formats, we developed a modular graphical user interface (GUI) built on the Python-native *pymmcore-plus* ecosystem (https://pymmcore-plus.github.io/pymmcore-plus/). While Micro-Manager offers broad hardware compatibility, its Java-based architecture limits workflow customization. Our GUI, *micromanager-gui* (https://github.com/fdrgsp/micromanager-gui), integrates *pymmcore-plus*, *pymmcore-widgets*, and *useq-schema* (https://pymmcore-plus.github.io/pymmcore-plus/) to enable synchronized stimulation, real-time segmentation, and standardized acquisition protocols within a modern, extensible Python framework. It remains fully compatible with existing Micro-Manager configurations, requiring no hardware or settings changes.

We extended the highly functional *MDAWidget* from the *pymmcore-widgets* library to create an intuitive interface for configuring multi-dimensional acquisitions, tailored to the needs of calcium imaging experiments. This widget supports key imaging modes including multi-channel, time-lapse, Z-stack, multi-position, and grid acquisition, while maintaining a user-friendly and flexible layout. Importantly, MDA settings can be easily saved and reloaded, enabling reproducibility of acquisition parameters across experiments. The widget also includes support for hardware autofocus devices, such as the Nikon Perfect Focus System (PFS) used in our experiments, which is essential for maintaining stable focus across wells during long or large-scale imaging sessions. To facilitate imaging across multi-well plates, we incorporated the *HCSWizard* feature from the *MDAWidget*, which provides a robust interface for configuring high-content screening workflows. This tool allows users to select from standard plate formats (e.g., 96- or 384-well) and perform precise plate calibration, with options to save, reload, and validate calibrations across sessions. It also supports flexible field-of-view (FOV) selection per well, offering three modes: acquisition at the well center, a reproducible random sampling of FOVs, or a defined grid layout within each well. This flexibility makes it easy to tailor acquisition strategies to different experimental needs while maintaining reproducibility and spatial consistency across datasets. For calcium imaging experiments, our preferred data format is OME-Zarr, which offers efficient, scalable storage well-suited to large, multidimensional datasets. The *MDAWidget* natively supports saving acquisitions in OME-Zarr, with Google TensorStore integrated as the backend for data writing. This allows chunked, asynchronous saving during acquisition, enabling large time-series datasets to be stored without requiring full in-memory representation.

### Data analysis

The *PlateViewer* GUI enables rapid data visualization and analysis at the level of individual positions or entire well plates, providing a timely overview of neuronal activity. Users can create customizable plate maps to annotate genotypes and treatment conditions, allowing for intuitive comparison of calcium activity across experimental groups. Annotated single-ROI data are automatically grouped by condition and FOV, and individual csv files summarizing each measured parameter are generated upon completion of the analysis.

### Compute ΔF/F_0_

After obtaining the segmentation file to delineate each neuronal soma and extracting the raw fluorescence traces over time for each ROI, the analysis proceeds with data pre-processing and ΔF/F₀ calculation. This step uses a sliding window approach combined with percentile-based background estimation. At each time point, the background fluorescence (F₀) is estimated by calculating the 10th percentile of fluorescence values within a user-defined temporal window. The ΔF/F₀ trace is then computed as the fractional change in fluorescence relative to this baseline (F - F₀) / F₀, and further baseline-corrected by subtracting its minimum value to ensure a zero-centered signal.

To extract the underlying neural activity from the calcium signals, we applied deconvolution to the ΔF/F₀ traces using the OASIS algorithm^40^. This method leverages an online active set strategy with an adaptive penalty parameter to account for the temporal dynamics of calcium indicators.

### Peak Detection

The next step in the analysis pipeline is the detection of calcium transients, which are identified as peaks in the deconvolved ΔF/F₀ traces. To achieve this, we employed the *scipy find_peaks* function, allowing for user-defined control over key detection parameters including prominence, minimum peak height, and minimum inter-peak distance. Noise levels for each ROI were estimated using the Median Absolute Deviation (MAD) method applied to the deconvolved traces. This involved computing the median, calculating the absolute deviations from the median, taking the median of those deviations, and normalizing the result by 0.6745 to approximate the standard deviation under the assumption of Gaussian noise (noise level = median(|dΔF/F₀-median(dΔF/F₀)|)/0.6745 where dΔF/F₀ is the deconvolved ΔF/F₀). The prominence threshold was then set adaptively for each ROI as the product of the noise estimate and a user-defined multiplier (default: 1.0), ensuring that detected peaks significantly exceeded local fluctuations. For the minimum height threshold, the software supports two modes. In the global mode, a fixed user-defined amplitude threshold is applied uniformly across all ROIs. In the adaptive mode, the height threshold is calculated for each ROI as a multiple of its noise level, allowing the detection algorithm to account for differences in signal-to-noise ratio across cells. Additionally, a minimum distance constraint between peaks (default: 2 frames) is enforced to prevent multiple detections of the same event, with this value configurable based on the imaging frame rate and calcium kinetics of the preparation.

### Features Extractions

Once peaks are detected, their amplitudes are extracted directly from the deconvolved ΔF/F₀ trace at the corresponding time points. In addition to peak amplitude, several basic features are computed for each ROI, including event frequency, inter-event interval (IEI), cell size (based on segmentation mask area) and percentage of active cells. Furthermore, population-level metrics such as synchrony and pairwise cross-correlation between ROIs are also calculated, providing insights into coordinated network activity. All thresholding parameters and extracted features are recorded to support downstream statistical and comparative analyses.

### Evoked Activity Experiments

For experiments involving optogenetic stimulation, the pipeline includes a dedicated evoked activity analysis module. Regions of interest (ROIs) are classified as “stimulated” if more than 10% of their area overlaps with a user-defined stimulation mask. Detected peaks are then classified as “stimulated” if they occur within a five-frame window following a stimulation pulse. A binary search algorithm is used to efficiently match peaks to stimulation events. Peak amplitudes are further grouped based on stimulation parameters, such as pulse duration and LED power. The system supports various forms of LED calibration (e.g., linear, quadratic, exponential, power-law, logarithmic), allowing for flexible power-response analysis.

### Visualization

At the multi-well level, the *PlateViewer* enables comparative analysis across wells grouped by genotype, treatment, or experimental condition. Summary statistics for amplitude, frequency, IEI, cell size, global synchrony, and activity rates are displayed using grouped bar plots with pooled standard errors. For evoked experiments, comparative metrics between stimulated and non-stimulated populations are also available, including power-response relationships and stimulation-dependent activity profiles. All visualizations dynamically update in response to changes in analysis parameters, offering an integrated and interactive environment for data interpretation. With real-time feedback, flexible filtering, and exportable figures, the *PlateViewer* supports both exploratory analysis and publication-quality visualization.

## Data and code availability

All data reported in this paper will be shared by the lead contact by request. All code is available on github (https://github.com/fdrgsp/micromanager-gui).

## CRediT authorship contribution statement

**WAS:** Writing – original draft, Visualization, Validation, Supervision, Methodology, Investigation, Formal analysis, Data curation, Conceptualization, Project administration, Funding acquisition. **FMG:** Writing – original draft, Visualization, Validation, Supervision, Methodology, Investigation, Formal analysis, Data curation, Conceptualization. **ZY:** Writing – review & editing, Investigation, Methodology, Validation, Data curation. **NAT:** Investigation, Data curation. **LL:** Investigation. **GS:** Investigation. **DW:** Methodology, Resources. **EDB:** Methodology, Resources. **KDW:** Methodology, Resources. **CC:** Investigation, Methodology, Resources. **MS:** Writing – review & editing, Supervision, Data curation, Project administration, Funding acquisition.

## Funding

This study was supported by the Rosamund Stone Zander Translational Neuroscience Center, Boston Children’s Hospital Equipment and Core Resources Allocation Committee award (to W.A.S.) and a grant from the Congressionally Directed Medical Research Program (W81XWH2110209 to M.S.).

## Acknowledgments

We would like to thank Dr Talley Lambert for his guidance. Dr Ivy Pin-Fang Chen, Ryan Chen, the Human Neuron Core at Boston Children’s Hospital (BCH IDDRC, P50HD105351) and the Viral Core at Boston Children’s Hospital (NIH5P30EY012196). We are grateful to members of the Sahin lab for critical reviewing of the manuscript. Finally, we would like to thank the patients and families who contributed to this work.

## Declaration of Competing Interest

Mustafa Sahin reports grant support from Novartis, Biogen, Astellas, Aeovian, Bridgebio, and Aucta. He has served on Scientific Advisory Boards for Novartis, Roche, Regenxbio, SpringWorks Therapeutics, Jaguar Therapeutics and Alkermes.

---

**Figure S1.**
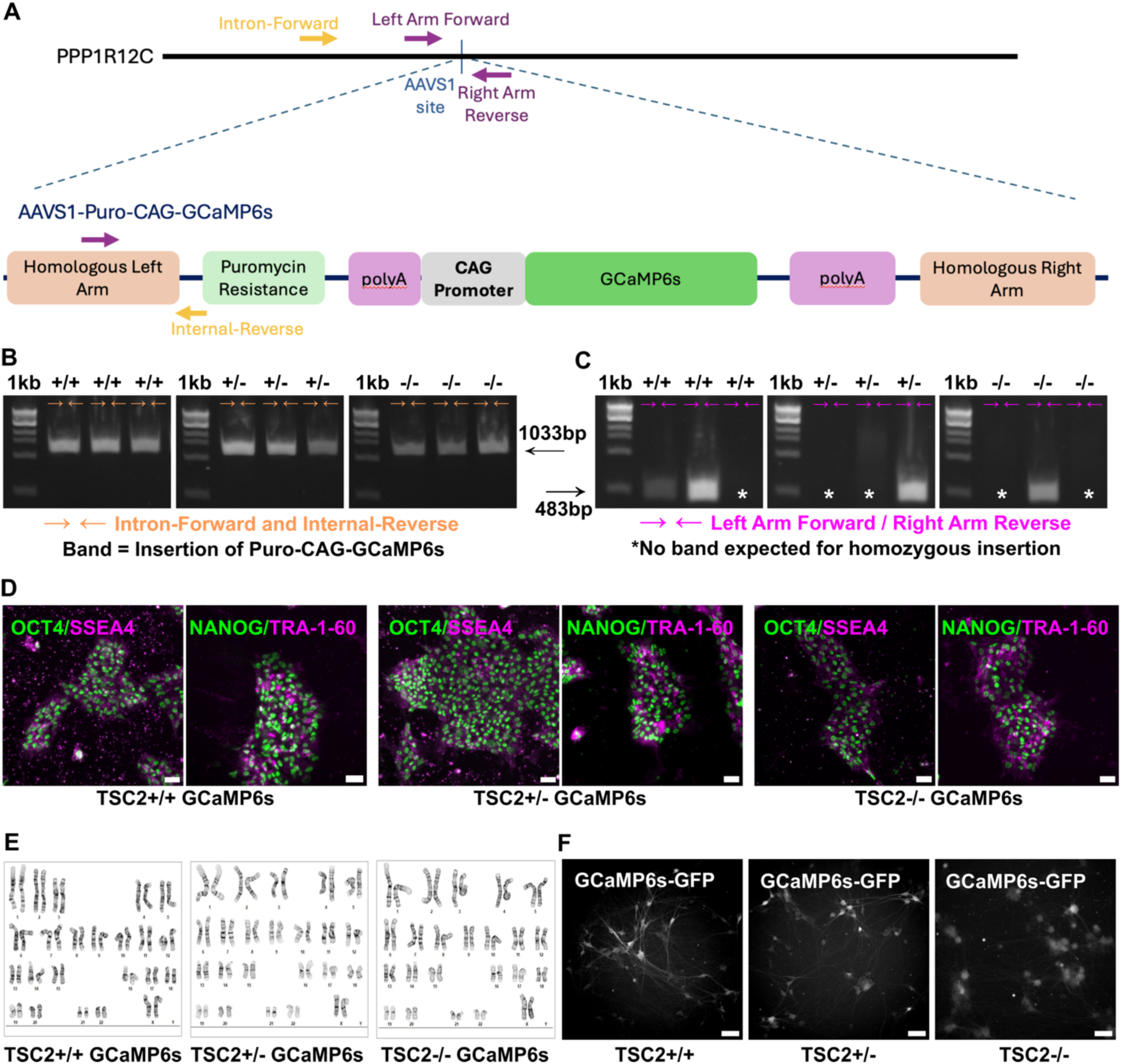
Generation of the TSC2 full allelic series GCaMP6s reporter lines. (A) Diagram of the strategy for the generation of the GCaMP6s reporter lines. (B) TSC2 full allelic series GCaMP6s reporter lines, primers: Intron-Forward and Internal-Reverse in orange on (A) and band size 1033bp. (C) TSC2 full allelic series GCaMP6s reporter lines, primers: Left Arm Forward and Right Arm Reverse in magenta in (A) and band size 483bp. No band expected for homozygous insertion, marked with a star (*). (D) Maintenance of pluripotency confirmed by expression of pluripotency markers: NANOG (Nanog homeobox x in green) and TRA-1-60 (podocalyxin in magenta), OCT4 (octamer binding transcription factor 4 in green) and SOX2 (SRY-Box Transcription Factor 2 in magenta), in undifferentiated pluripotent hiPSC colonies for the TSC2 full allelic series after insertion of GCaMP6s. (E) G-banded karyotype for the TSC2 full allelic series post-gene editing showing a normal karyotype for all lines (F) GCaMP6s-GFP expression following differentiation of the TSC2 full allelic hiPSCs series, at DIV42, scale bar 100µm.

**Table S1:**
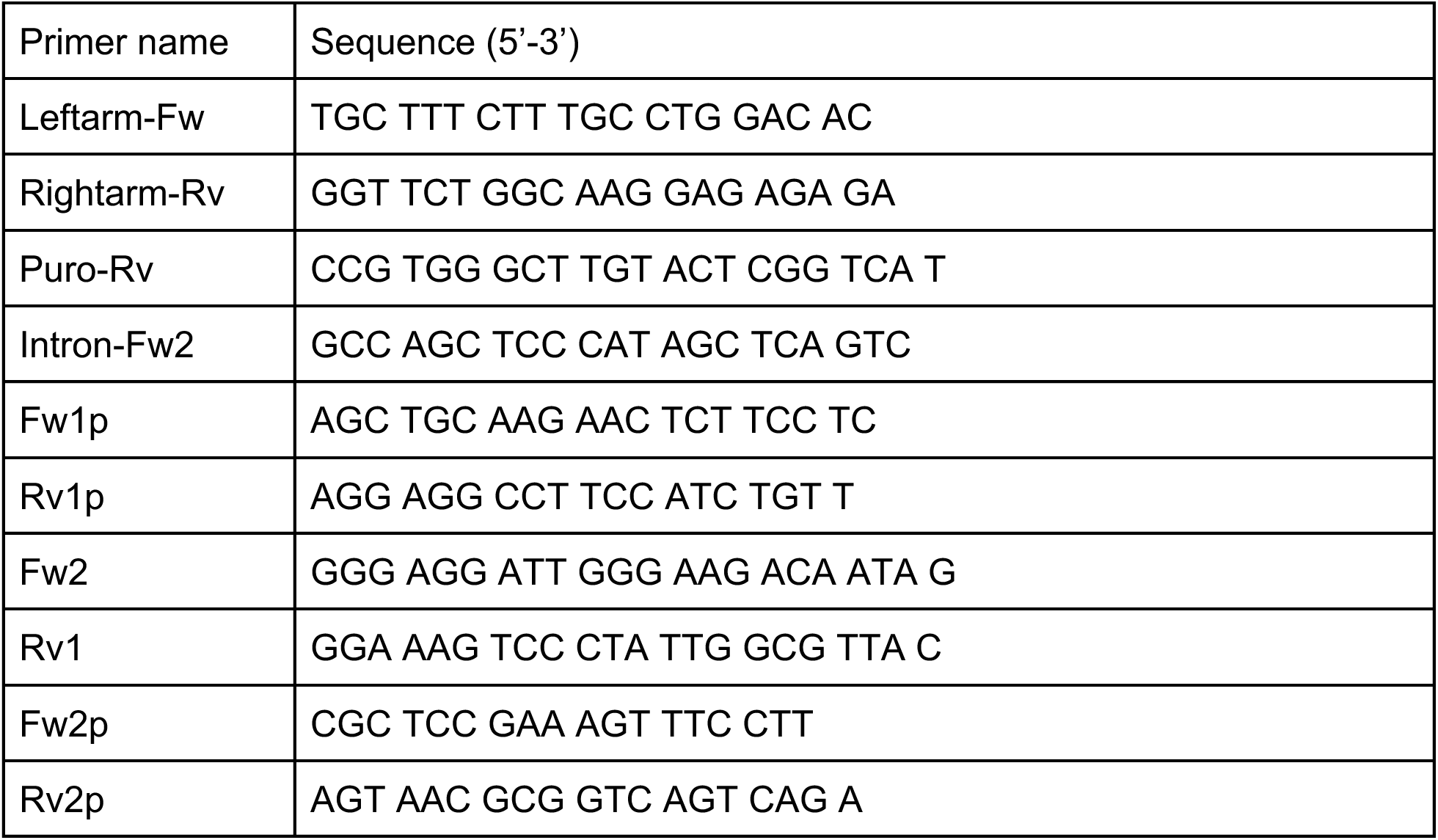
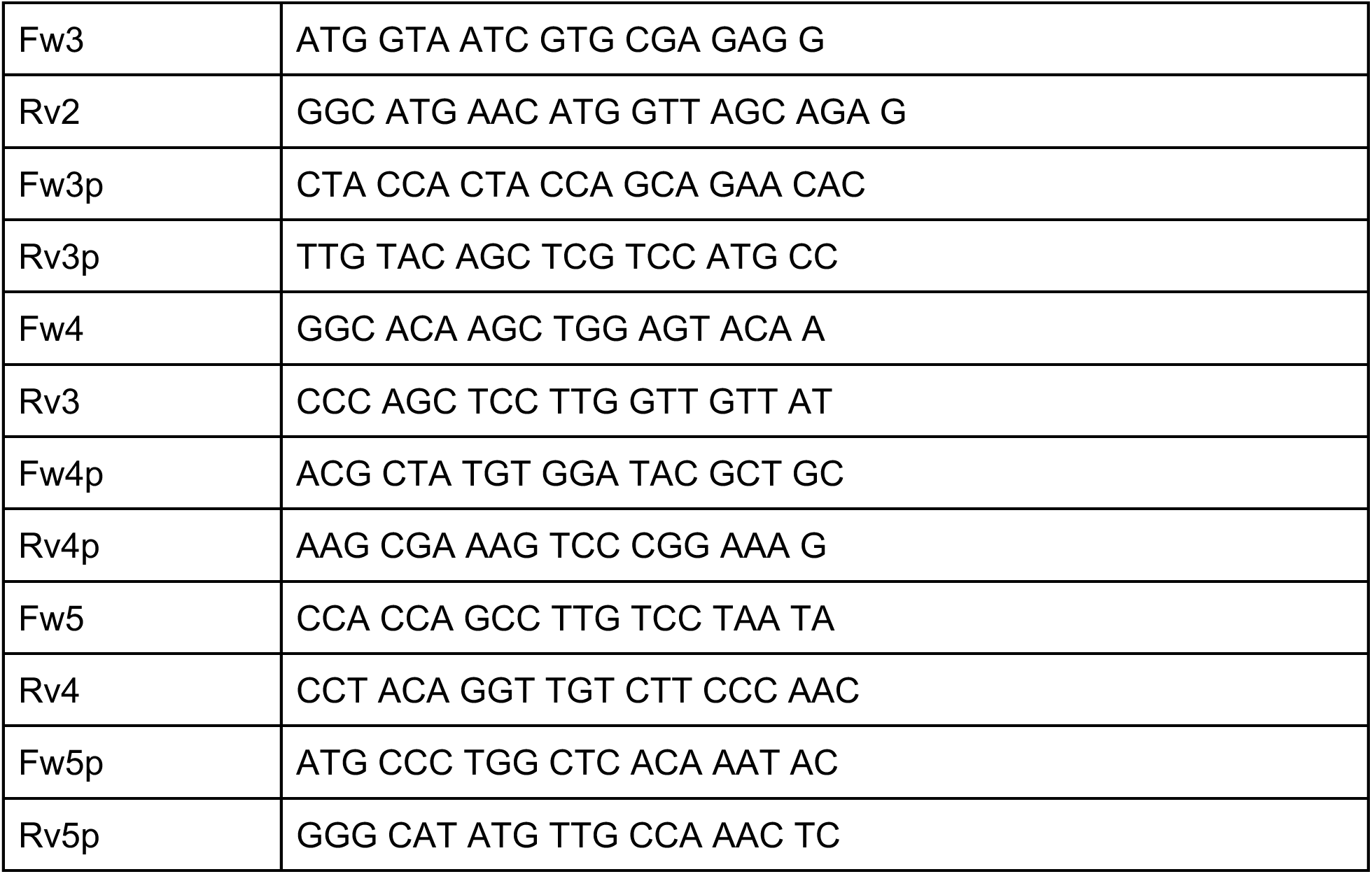
Primer sequences to confirm the plasmid insertion (AAVS1-Puro-CAG-GCaMP6s)

**Table S2:** Antibodies used for pluripotency marker staining.

| Staining | Antibody | Vendor | Catalog # | Host | Dilution |
| --- | --- | --- | --- | --- | --- |
| Primary antibody | Anti-OCT4 | Cell Signaling | 2840s | Rabbit | 1:250 |
|  | Anti-SSE4 | Invitrogen | 41-4000 | Mouse | 1:250 |
|  | Anti-Tra1-60 | Invitrogen | 41-1000 | Mouse | 1:250 |
|  | Anti-NANOG | Abcam | AB109250 | Rabbit | 1:250 |
| Secondary antibody | Anti-Rabbit Alexa 647 | Invitrogen | A21245 | Goat | 1:500 |
|  | Anti-Mouse Alexa 568 | Invitrogen | A11004 | Goat | 1:500 |

## Notes

https://github.com/fdrgsp/micromanager-gui

