## Supplementary figures and images for "An open-source pipeline for calcium imaging and all-optical physiology in human stem cell-derived neurons"

### Supplementary Figure 1

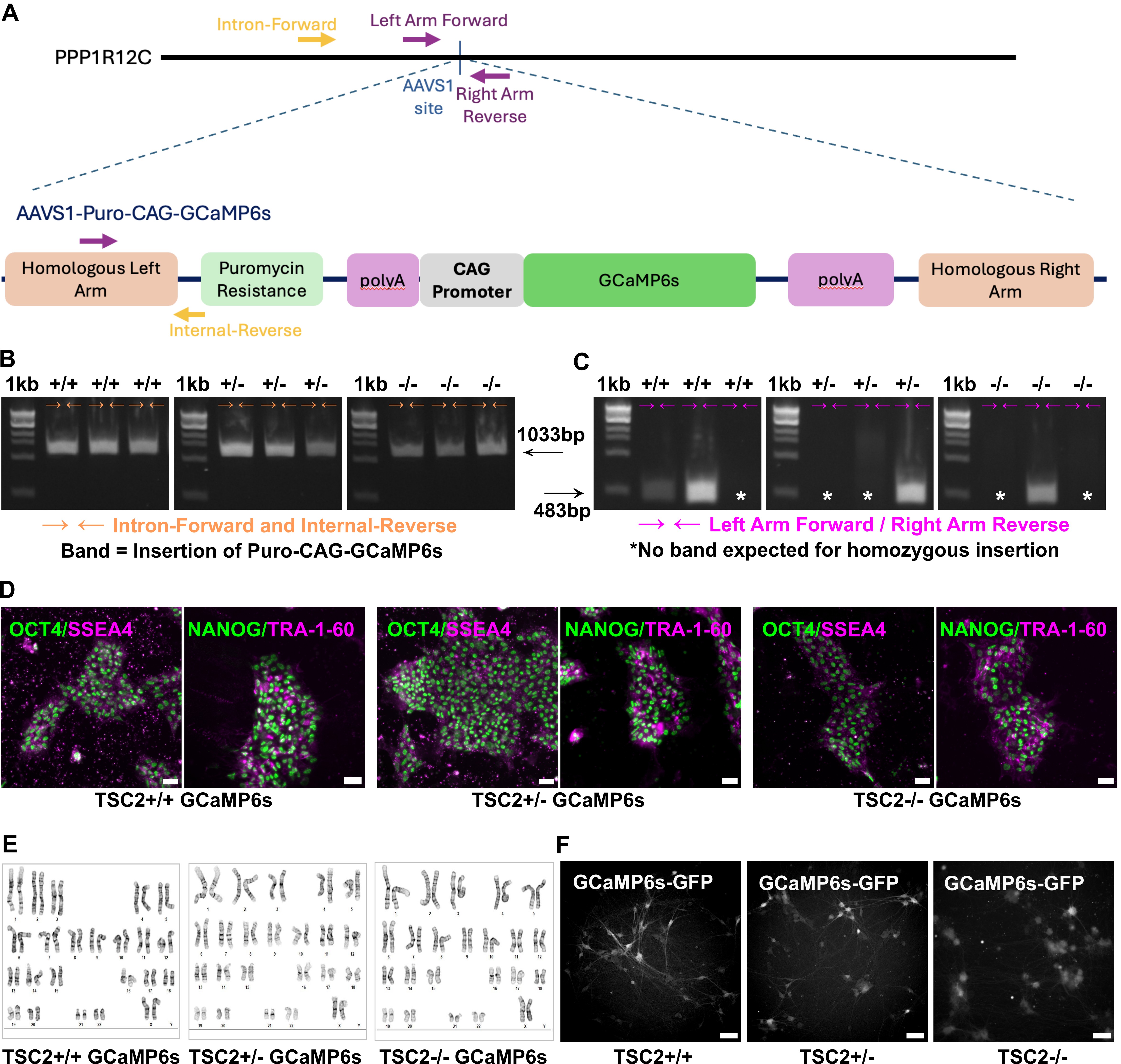
